# Membrane insertion function for SUN-KASH complex revealed by high resolution analysis of yeast centrosomes

**DOI:** 10.1101/413799

**Authors:** Jingjing Chen, Jennifer M. Gardner, Zulin Yu, Sarah E. Smith, Sean McKinney, Brian D. Slaughter, Jay R. Unruh, Sue L. Jaspersen

## Abstract

Bipolar spindle formation in yeast requires insertion of centrosomes (known as spindle pole bodies (SPBs)) into fenestrated regions of the nuclear envelope (NE). Using structured-illumination microscopy and bimolecular fluorescence complementation, we map protein distribution at SPB fenestra and interrogate protein-protein interactions with high spatial resolution. We find that the Sad1-UNC-84 (SUN) protein Mps3 forms a ring-like structure around the SPB, similar to toroids seen for components of the SPB insertion network (SPIN). Mps3 and the SPIN component Mps2 (a Klarsicht-ANC-1-Syne-1 domain (KASH)-like protein) form a novel non-canonical linker of nucleoskeleton and cytoskeleton (LINC) complex that is connected in both luminal and extraluminal domains. This hairpin-like LINC complex forms during SPB insertion, suggesting it functions in NE reorganization at the pore membrane. The LINC complex also controls the distribution of a soluble SPIN component Bbp1. Taken together our work shows that Mps3 is a fifth SPIN component and suggests both direct and indirect roles for the LINC complex in NE remodeling.

## Introduction

The double lipid bilayer of the nuclear membrane serves as a physical barrier to restrict movement of macromolecules from the cytoplasm to nucleus, or vice versa. Throughout interphase, transport across the nuclear envelope (NE) is facilitated by nuclear pore complexes (NPCs) that are located at sites where the inner and outer nuclear membranes (INM and ONMs) are contiguous, known as the pore membrane. In fungi, as well as in rapidly dividing cells such as Drosophila and C. *elegans* embryos, the INM and ONM also come together to form a fenestra at or near the microtubule-organizing center (MTOC) (reviewed in (Funakoshi et al., 2011; Smoyer and Jaspersen, 2014)), which is known as the centrosome in metazoans and the spindle pole body (SPB) in yeast. Unlike most metazoans, the NE does not breakdown in these systems during mitosis, so integration of the SPB into the NE in yeast, for example, ensures that microtubules can form a mitotic spindle to segregate the genome within the nucleus while simultaneously nucleating cytoplasmic microtubules that orient the nucleus for delivery of a genome into each of the daughter cells.

In budding yeast, the SPB is anchored in a fenestrated region of the NE throughout the cell cycle (reviewed in (Cavanaugh and Jaspersen, 2017; Ruthnick and Schiebel, 2016)). Genetic analysis suggests that SPB incorporation into the NE requires at least four factors: a soluble SPB protein, Bbp1; an amphipathic domain-containing protein Nbp1; the dual SPB-NPC transmembrane protein Ndc1; and a Klarsicht-ANC-1-Snye-1 homology (KASH)-like protein Mps2 (Araki et al., 2006; Chial et al., 1998; Munoz-Centeno et al., 1999; Schramm et al., 2000; Winey et al., 1991). Known as the SPB insertion network (SPIN) (Ruthnick et al., 2017), these components display extensive genetic and physical interactions and are thought to form a donut-like structure around the core SPB, suggesting roles for the SPIN in both NE fenestration and SPB anchorage in the NE. Interestingly, specific NPC components genetically interact with the SPIN, leading to the idea that NPCs and SPBs share common regulators or insertion factors,including NPC components themselves (Casey et al., 2012; Chen et al., 2014; Chial et al., 1998; Lau et al., 2004; Ruthnick et al., 2017; Sezen et al., 2009; Witkin et al., 2010). How SPIN components or NPCs lead to NE fenestration is not understood, but data in both mammals and yeast suggest that Sad1-UNC-84 (SUN) domain proteins also may be involved in this process (Bestul et al., 2017; Fernandez-Alvarez et al., 2016; Friederichs et al., 2011; Talamas and Hetzer, 2011).

Central to understanding how complexes such as the SPB and NPC are assembled and anchored in the membrane is the need to develop rigorous, reproducible methods to compare NE-associated protein structures at high resolution. Here, we describe how structured-illumination microscopy (SIM), iterative three-dimensional single particle averaging (SPA) and bimolecular fluorescence complementation (BiFC) can be combined to study the organization of SPIN proteins at the SPB in wild-type and mutant cells. This approach led to the surprising discovery that the SPIN forms at least two domains, one that contains Bbp1 and a Bbp1-independent region. We show this is due, at least in part, to the budding yeast SUN protein Mps3, which forms an a-typical SUN-KASH complex with Mps2.

## Results and Discussion

### Radial distribution of SPIN components using 3D particle averaging

To understand the role of the SPIN in NE fenestration, we were interested in SPIN protein distribution in wild-type cells. The SPIN components Ndc1 and Mps2 were observed to surround the SPB core (Spc42) using SIM, a localization pattern that we refer to as a ring or toroid (Fig 1A-B) (Burns et al., 2015; Ruthnick et al., 2017). Nbp1 and Bbp1 only appeared as rings in a small fraction of asynchronously growing cells (Burns et al., 2015) or in synchronized cells (Ruthnick et al., 2017) in previous work, suggesting a transient toroidal localization during SPB insertion in late G1. However, as some of these experiments were conducted in cells overproducing a Bbp1-interacting protein, it is possible that Bbp1 is artificially recruited to the toroid. Therefore, we created diploid strains with endogenously mTurquoise2 (mT2)-tagged SPIN components and used SIM in asynchronously growing cells to examine localization relative to the SPB toroid (Ndc1-YFP) and core (Spc42-mCherry). Mps2-mT2 and Nbp1-mT2 colocalized with Ndc1-YFP at the SPB toroid in both individual images (Fig 1B) and in averaged toroids, which were generated from multiple, randomly oriented SPBs that were computationally aligned, reconstructed and normalized using Ndc1-YFP (Fig 1C-D; Fig S1A-B). These experiments showed that the size of Ndc1-YFP and Mps2-mT2 toroids are similar to each other (169±1 nm and 170±1 nm, respectively) and to estimates of SPB diameter (160 nm in diploids) determined by EM (Byers and Goetsch, 1974; Li et al., 2006), validating our re-alignment and normalization protocol. In contrast, the Nbp1 toroid diameter (135±1 nm) was 20% smaller (Fig 1E), perhaps explaining the difficulty in visualizing Nbp1 as a toroid in earlier work using the longer-wavelength YFP fluorophore (Burns et al., 2015).

**Figure 1.**
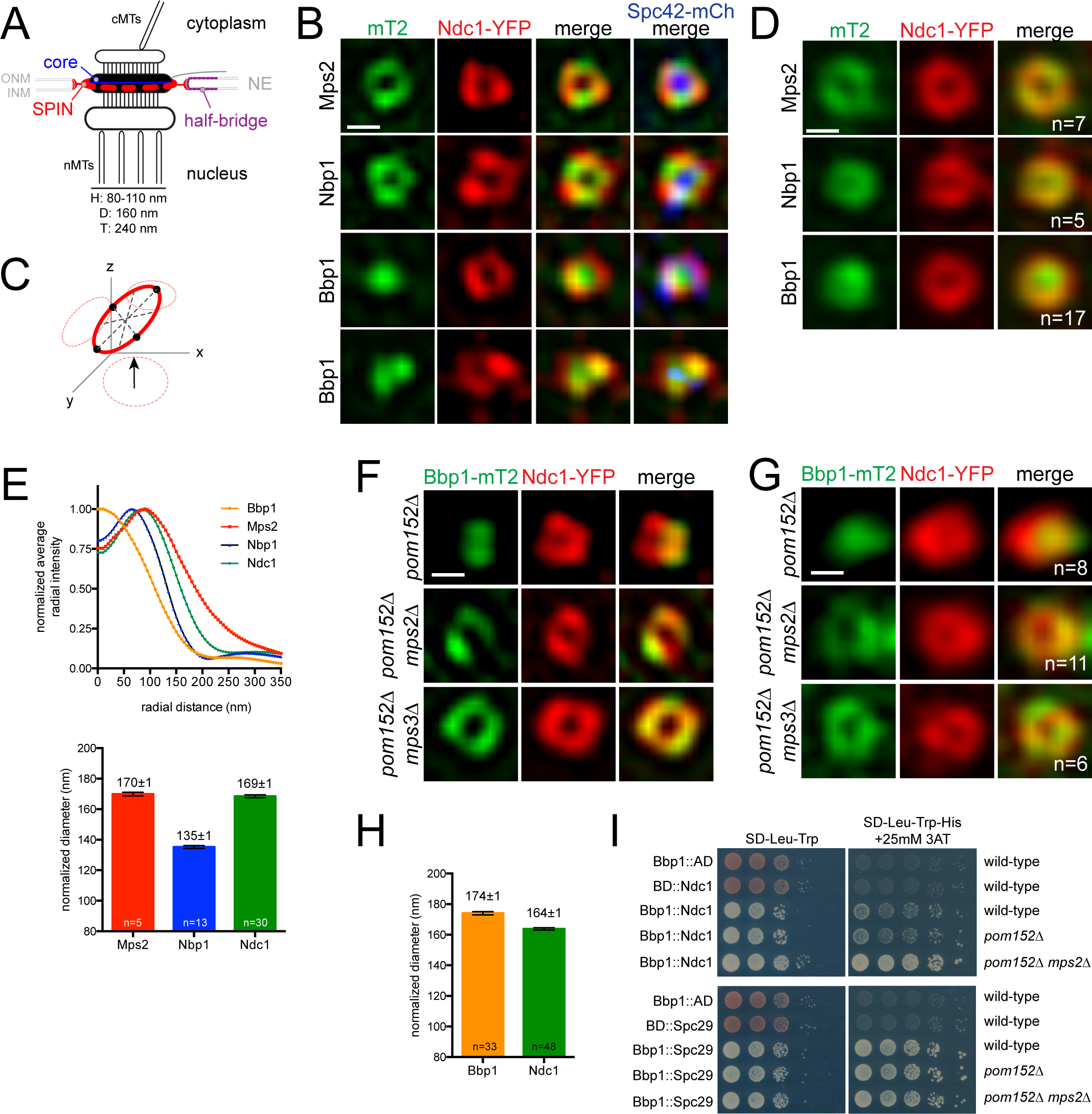
Radial distribution of SPIN components at the SPB. A. Schematic of the SPB showing how the soluble core is thought to be tethered to the NE by the SPIN. The membrane-associated bridge is also shown. SPB dimensions in haploid (H), diploid (D) or tetraploid (T) cells based on EM measurements are listed (Adams and Kilmartin, 2000; Byers and Goetsch, 1975). B. Representative SIM images with top down view of SPB. Cells contain Ndc1-YFP (red) and Spc42-mCherry (blue) to detect the SPB ring and core, respectively, along with the indicated protein tagged with mT2 (green) (SLJ11171, Mps2; SLJ10898, Nbp1; SLJ10635, Bbp1). C-D. As shown in the schematic (C), averaged images (D) were generated by realigning multiple SPB rings three-dimensions (see Fig S1A-B). Number of averaged rings is shown. E. Fluorescence profiles of SPIN components from the center of the SPB outwards, based on the projections in (D). Average ring diameter was determined in aligned images based on the center of Gaussian fits of fluorescence intensity. Because Ndc1-YFP diameter varied by up to 20 nm between different strain isolates, values were normalized using Ndc1-YFP values. Error bars, SEM based on the number of points shown. P values were calculated using the students t-test; only Nbp1 was statistically significant. F. SIM showing localization of Ndc1-YFP (red) and Bbp1-mT2 (green) in asynchronously grown *pom152Δ* (SLJ12302), *pom152Δ mps2Δ* (SLJ10998) and *pom152Δ mps3Δ* (SLJ10534) strains. G. Averaged images were generated by re-aligning multiple SPB rings three-dimensions, as in (C). Number of averaged rings is shown. H. Average ring diameter was determined in aligned images based on the center of Gaussian fits of fluorescence intensity as in (E). I. Pairwise protein interactions between Bbp1 fused to the *GAL4*-binding domain (BD) and Ndc1 or Spc29 fused to the *GAL4*-activation domain (AD) were tested by serial dilution assays in the yeast two-hybrid system in wild-type (SLJ1644), *pom152Δ* (SLJ12623) and *pom152Δ mps2Δ* (SLJ12624). As a control, empty BD and AD vectors were also used. Growth on media lacking tryptophan (Trp), leucine (Leu) and histidine (His) that also contained 25 mM 3-AT (right) indicates an interaction, while growth on -Trp-Leu is a plating control. Bars in Fig 1, 100 nm.

Unlike other SPIN components, Bbp1-mT2 typically did not localize to a toroid but rather formed one or two large puncta (Fig 1B). In rare instances (7/63), a ring-like distribution of Bbp1-mT2 was detected in unbudded G1, medium-budded S phase and large-budded mitotic cells, making it unlikely that Bbp1-mT2 toroidal distribution is cell cycle regulated. Therefore, we considered the possibility that the distribution of Bbp1-mT2 was spatially controlled by its primary binding partner, Mps2 (Kupke et al., 2017; Schramm et al., 2000). *MPS2* can be deleted in yeast strains lacking the nucleoporins *POM34* or *POM152* (Katta et al., 2015; Kupke et al., 2011; Witkin et al., 2010). In the absence of *MPS2*, Bbp1-mT2 distributed to ring-like structures that co-localize with Ndc1-YFP (Fig 1F-G; S1C-D). The size of the Bbp1-mT2 toroid (174±1 nm) was similar to that of Ndc1 and Mps2 (Fig 1H), suggesting that loss of *MPS2* might allow Bbp1 to bind to another SPIN or SPB component at the membrane region surrounding the SPB core. Consistent with this idea, we found that Bbp1 binding to Ndc1 increased in cells lacking *MPS2* (Fig 1I). Taken together, these data support the idea that the SPIN components Ndc1, Mps2 and Nbp1 constitutively localize to a ring-like structure surrounding the SPB in wild-type cells. Unexpectedly, our data show that the SPIN toroid has at least two domains – a region that includes Mps2, Nbp1 and Ndc1 and a second that also contains Bbp1.

### Mps3 is a component of the SPB toroid and bridge

Mps3 is a highly divergent SUN domain-containing protein that binds to the C-terminus of Mps2 in the luminal space forming a linker of nucleoskeleton and cytoskeleton (LINC) bridge across the INM and ONM (Jaspersen et al., 2006). Deletion of *MPS3*, like *MPS2*, results in redistribution of Bbp1-mT2 to a toroid (Fig 1F-G), suggesting that Mps3, at least indirectly, affects SPB fenestrae. Consistent with this idea, a dominant allele of *MPS3* has defects in SPB insertion similar to other SPIN mutants (Friederichs et al., 2011) and, if present in molar excess, Mps3 can interact with a Mps2-Bbp1 complex in vitro (Kupke et al., 2017).

To determine if Mps3 shows a toroidal distribution, we examined Mps3-mT2 localization by SIM in a diploid strain containing Ndc1-YFP and Spc42-mCherry. In individual and merged images (Fig 2A-B), Mps3-mT2 surrounded the SPB similar to the ring-like distribution recently described for the fission yeast SUN protein Sad1 (Bestul et al., 2017). It is unclear why Mps3 was not observed in a toroid in a recent study (Ruthnick et al., 2017), but it may relate to the use of overexpressed *SPC42* and *SPC29*, which alter SPB architecture (Donaldson and Kilmartin, 1996; Schramm et al., 2000). Mps3’s distribution was different than other SPIN components in that a significant fraction of Mps3-mT2 also appeared as a large focus on one side of the SPB, which corresponded to the bridge based on co-localization with YFP-Sfi1, a cytoplasmic bridge protein (Fig 2D). Both rings and bridge-localized Mps3 can be seen in duplicated side-by-side SPBs, suggesting that the toroidal distribution of Mps3 is linked to SPB insertion (Fig 2D). Alignment and normalization of toroids revealed that Mps3 diameter in the x direction that does not include the bridge localized population is 167±3 nm while the length in the y axis is 194±1 nm (Fig 2C).

**Figure 2.**
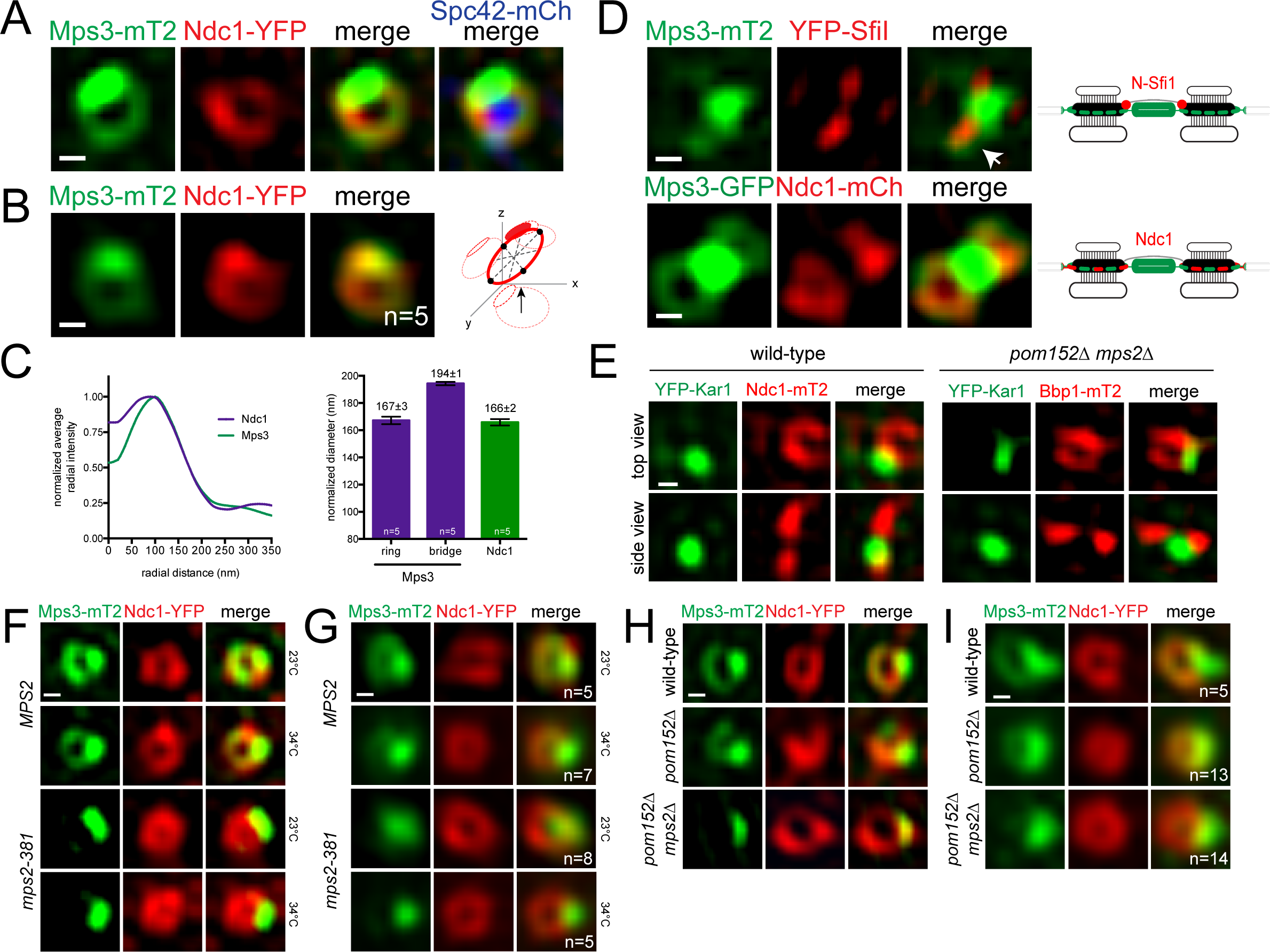
Mps3 localization to a toroid surrounding the SPB is Mps2-dependent. A. Representative SIM image from cells containing Ndc1-YFP (red), Spc42-mCherry (blue) and Mps3-mT2 (green) (SLJ10636). B. Averaged images from the indicated number of rings from A. The asymmetric distribution of Mps3-mT2 facilitated realignment. C. Average ring diameter was determined in aligned images based on Gaussian fits of fluorescence intensity, using Ndc1-YFP to normalize measurements. Because Mps3-mT2 was anisotropic, its diameter varied based on the region selected for analysis; shown are the diameters from the ring region only and from the region that includes both the ring and the half-bridge. Error bars, SEM based on the number of points shown. D. In G1 cells containing YFP-Sfi1 (red) and Mps3-mT2 (green) (SLJ12060), Mps3 is present between the Sfi1 foci that mark the ends of the extended bridge and in a ring (arrow). In SLJ5496 cells released from α-factor for 40 min, Mps3-GFP (green) co-localizes with Ndc1-mCh (red) at toroids. Schematics illustrate protein distribution at SPBs. E. YFP-Kar1 distribution in SIM images taken to show the SPB from a top-down and side-on view from wild-type (SLJ10001) and *pom152Δ mps2Δ* (SLJ12620) strains. F. SIM showing distribution of Mps3-mT2 (green) along with Ndc1-YFP (red) in wild-type (SLJ12772) and *mps2-381* (SLJ12616) cells grown at 23°C or shifted to 34°C for 3 h.G. Averaged images were generated by re-aligning the indicated number of SPB rings three-dimensions.H-I. Individual SIM (H) and averaged (I) images showing localization of Ndc1-YFP (red) and the distribution of Mps3-mT2 (green) in wild-type (SLJ10636), *pom152Δ* (SLJ11071), *pom152Δ mps2Δ* (SLJ10535). Bars in Fig 2, 100 nm.

Toroid formation is not a general feature of bridge components as we did not detect other soluble or membrane proteins such as Sfi1 or Kar1 surrounding the SPB core (Fig 2E). Thus, Mps3 uniquely exists in three populations: the INM (Fig S2A), the SPB bridge and the toroid surrounding the core SPB. Quantitation of the average realigned fluorescence distribution indicates that roughly half (45±5%, n=6) of Mps3 protein at the SPB is located in the toroid, with the remainder (55±5%, n=6) densely packed in the bridge region extending away from the core SPB. Based on its distribution at the toroid, effect on Bbp1 localization, role in SPB insertion and Mps2 binding, we propose that Mps3 is a novel component of the SPIN.

### Mps3 toroid formation is Mps2-dependent

The idea that the SPB membrane is separated into subdomains raises the interesting question as to how a protein such as Mps3 is localized to distinct areas such as the toroid and bridge. Previous genetic and binding data using *mps2-381*, a C-terminal truncation, showed that Mps2 and Mps3 interact through a luminal linkage reminiscent of other KASH and SUN proteins. This interaction was proposed to tether the bridge to the core SPB (Jaspersen et al., 2006), but it is possible that Mps2 binding recruits Mps3 to the toroid in addition to, or instead of, the bridge.

Examination of Mps3-mT2 distribution in *mps2-381* showed Mps3 was lost specifically from the toroid in the *mps2-381* mutant at both 23° and 34°C (Fig 2F-G), suggesting that Mps2 plays a role in Mps3 distribution at the ring but not the bridge. To exclude the possibility that residual binding to mps2-381 tethers Mps3 to the bridge (Jaspersen et al., 2006), we also showed that Mps3-mT2 is absent from toroids but present at the bridge and INM in *pom152Δ mps2Δ* and *pom34Δ mps2Δ* cells (Fig 2H-I; S2A-C). This loss is not due to *pom152Δ* or *pom34Δ* nor is it caused by gross structural abnormalities at the SPB, as other SPIN components such as Ndc1 and Npb1 localized to the toroid and the laminar structure of the SPB in *pom152Δ mps2Δ* was morphologically indistinguishable from wild-type by EM (Fig S2D-G). Deletion of *MPS3* together with either *POM34* or *POM15*2 had no effect on Mps2-mT2 and Nbp1-mT2 distribution, as shown in individual or averaged (Fig S2F-I) images. Thus, at least two forms of Mps3 exist at the SPB —a toroid-specific population of Mps3 that requires Mps2 for its formation and/or stabilization, and a second population that localizes to the bridge independently of interaction with Mps2. Our finding that SUN protein (Mps3) localization is dependent on the KASH-like protein (Mps2) but Mps2 localization is Mps3-independent is distinct from most other SUN-KASH interactions.

### Mps3 binds to Mps2 throughout the toroid

Using fluorescence resonance energy transfer (FRET), we assayed Mps3’s interactions with other SPIN components, taking into account the relative abundance of donor and acceptor proteins and protein topology, which both affect FRET (Fig 3A-C). We did not detect FRET between YFP-Mps3 and Ndc1-mT2 (-5.7±1.5%, n=213), and our FRET between YFP-Mps3 and Bbp1-mT2 (2.6±1.3%, n=224, p=0.16) was not statistically significant compared to controls (Fig 3D-E). However, FRET between YFP-Mps3 and Nbp1-mT2 was 5.7±0.7% (n=795), similar to FRET levels observed between other SPIN components (Fig 3D-E). We were unable to test FRET between the N-termini of Mps2 and Mps3 because the tagged strains were lethal in combination. Strikingly, the 39.9±2.1% (n=271) FRET between the C-termini of Mps3 and Mps2 was more than double that of any other protein pair examined, including our positive FRET control (Fig 3D-E). This very high FRET indicates that multiple copies of the Mps2 C-terminus interact with a single Mps3 C-terminus, suggesting an alternative high stoichiometry complex compared to the SUN2-KASH1/2 trimer (Sosa et al., 2012). While this is perhaps not surprising given that Mps3 lacks key residues that mediate the SUN-KASH interface (Sosa et al., 2012), it raises the possibility that SUN proteins, particularly those involved in centrosome tethering (Meier, 2016), may interact with KASH-like proteins using alternative mechanisms.

**Figure 3.**
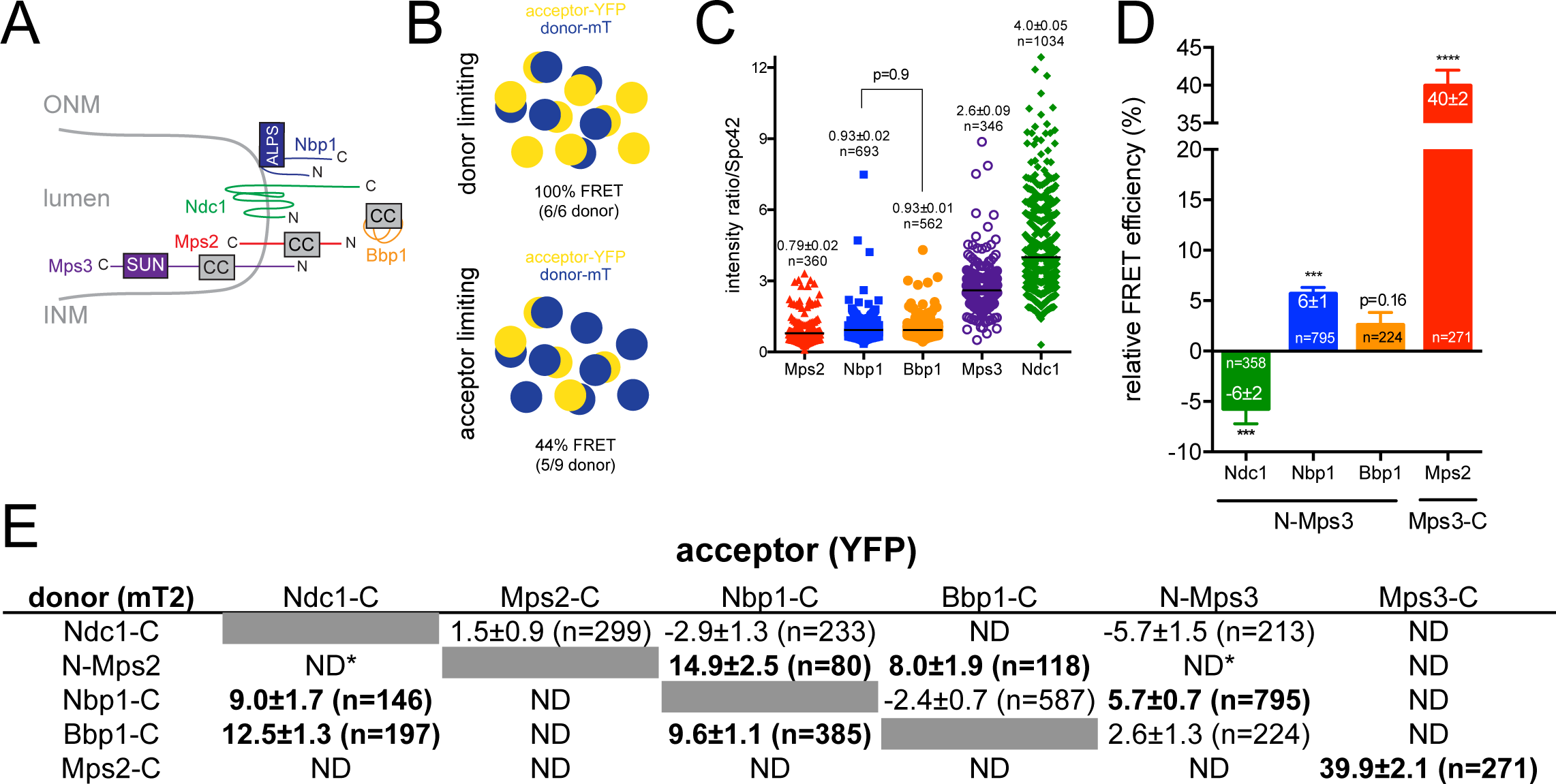
Mps3 binds to SPIN components at the toroid. A. Schematic showing the contiguous INM and ONM at the SPB and the topology of Ndc1, Mps2 and Mps3. Nbp1 is not integral to the membrane but is thought to interact with a leaflet via its amphipathic helix (Kupke et al., 2011). Bbp1 is soluble (Schramm et al., 2000). Also depicted is the location of the Mps3 SUN domain and the location of one or more coiled-coils (CC) in SPIN components. B. Acceptor photo-bleaching FRET is sensitive to protein abundance. Note that this simplified model assumes that all molecules are capable of FRET, which is unlikely to be the case for Mps3. It also does not account for conformational changes that may affect FRET efficiency. Adapted from (Katta et al., 2015). C. Levels of Mps2- (SLJ8065), Nbp1-(SLJ12020), Bbp1-(SLJ11903), Mps3-(SLJ8835) and Ndc1-mT2 (SLJ7941) in asynchronously growing haploid cells at the SPB were determined relative to the amount of Spc42-YFP. Long bars depict average values, which are listed with SEM based on the number of points shown. P values were calculated using Student’s t-test; all are significant (p<0.0001) except for Nbp1-mT2 and Bbp1-mT2. D. Binding between Mps3 and SPIN components was analyzed at the SPB using acceptor photo-bleaching FRET in asynchronously growing cells. Average FRET efficiency in the number of cells analyzed is listed along with SEM. Negative FRET values are most likely due to bleaching of the donor since we excluded cells in which the SPBs moved (see Materials & Methods). P values were determined using the Student’s t-test compared to the donor only control. For comparison, we observed 9.1±0.7 (n=398) and -1.4±0.5 (n=484) percent FRET in the positive (Spc42-mT2/Cnm67-YFP, SLJ8173) and negative (YFP-Spc110-mT2, SLJ7987) controls, respectively (Katta et al., 2015; Muller et al., 2005). E. Acceptor photo-bleaching FRET between SPIN components and with Mps3. Average FRET efficiency is listed along with SEM; the number of cells analyzed is listed. Some FRET pairs were not examined because of protein topology (ND) or incompatibility between tagged proteins (ND*).

To look at the spatial localization of discrete protein-protein interactions within a large complex, such as Mps2 binding to Mps3 at the bridge or toroid, we combined BiFC with SIM. Reconstituted GFP (rGFP) was observed at the toroid in strains containing GFP_11_-mCherry-Mps2 and Ndc1-GFP_1-10_, Nbp1-GFP_1-10_ or GFP_1-10_-Mps3 (Fig 4B-D). A variety of controls showed that this signal was specific and biologically relevant, including analysis of Bbp1-GFP_1-10_, which did not interact with GFP_11_-mCherry-Mps2 throughout the toroid but formed a specific rGFP focus adjacent to the bridge (Fig 4B). Although Mps3 and Bbp1 both bind to Mps2 (Jaspersen et al., 2006; Kupke et al., 2017; Schramm et al., 2000), our data show that the interactions are spatially distinct, with Mps2-Mps3 complexes around the toroid and more restricted Mps2-Bbp1 complexes (Fig 4F). That Bbp1 is able to localize to the toroid in cells lacking *MPS3* suggests that Mps3 and Bbp1 organization are coordinated, possibly through competition for the same binding site on Mps2 or other temporal regulation.

**Figure 4.**
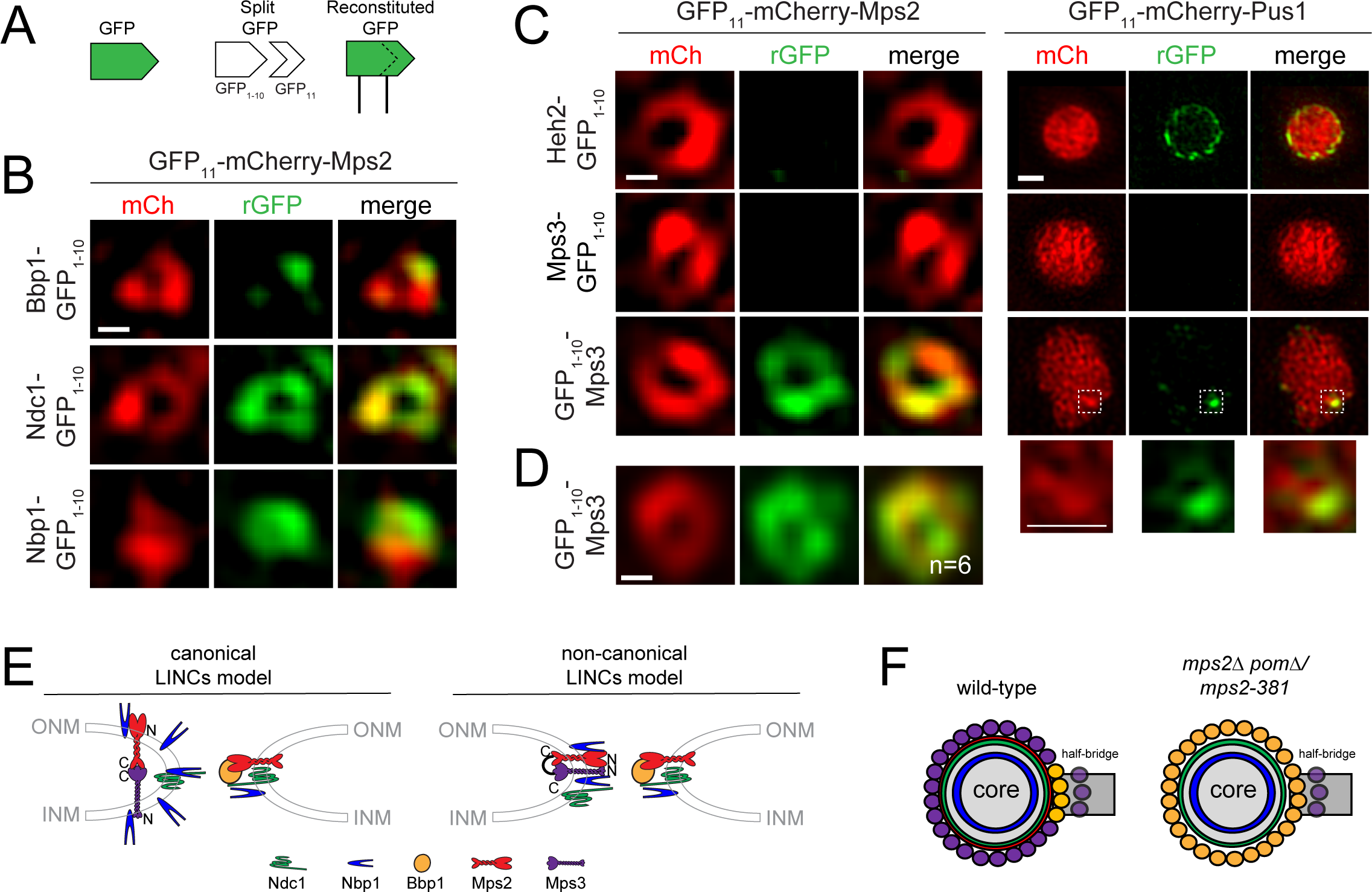
BiFC combined with super resolution reveals a non-canonical LINC. A. GFP can be split into two non-fluorescent halves, GFP_1-10_ and GFP_11_, which can reconstitute fluorescence if present in the same subcellular compartment (Cabantous and Waldo, 2006). B. SIM from cells expressing GFP_11_- mCherry-Mps2 (red) together with Ndc1-GFP_1-10_ (SLJ12825), Nbp1-GFP_1-10_ (SLJ12826), Bbp1-GFP_1-10_ (SLJ12866) to detect rGFP (green) at the SPB. C. SIM from cells expressing the soluble nuclear reporter GFP_11_-mCh-Pus1 or GFP_11_-mCherry-Mps2 (red) to detect interactions at the NE or SPB, respectively, via rGFP (green) with Heh2-GFP_1-10_ (SLJ8138/12773), Mps3-GFP_1-10_ (SLJ9399/12476) or GFP_1-10_-Mps3 (SLJ8577/12474/). Although GFP_11_-mCh-Pus1 is present through the nucleoplasm, reconstituted GFP at the SPB can be seen with GFP_1-10_-Mps3 but not with other samples. A magnified view of the SPB region shows that rGFP is highly asymmetric, similar to GFP localization of Mps3, indicating that both the toroid and half-bridge are accessible to GFP_11_. D. Averaged images of rGFP rings present in GFP_1-10_-Mps3/GFP_11_- mCherry-Mps2 cells were generated by re-aligning the indicated number of cells in using mCh in three-dimensions. E-F. Based on the FRET data in (Fig 3) and BiFC/SIM data in (Fig 4B-D), interactions between SPIN components can be summarized in models in which the N-termini of Mps2 and Mps3 are located in the cytoplasm and nucleoplasm, respectively (canonical LINCs model), or they are located at the pore membrane (non-canonical LINCs model). In both cases, the C-termini of Mps2 and Mps3 interact within the luminal space. Our data also suggest the presence of at least two separate membrane domains, distinguished by the presence (right) or absence of Bbp1 (left) (wild-type, F). Mps2-Mps3 binding appears to occur at the toroid, rather than the half-bridge, suggesting two forms of Mps3 also exist (F). It is unknown what tethers Mps3 to the half-bridge. Although Kar1 has been proposed to be a KASH-like protein, it contains a single amino acid in the luminal space. Bars in Fig 4, 100 nm, 2 µm in Figure 4C (right panel).

Rather than forming a canonical LINC complex (Fig 4E, left), our BiFC data showing that N-termini interact suggests that the LINC complex folds back on itself in a hairpin at the pore membrane (Fig 4E, right). This model is attractive in that Nbp1 is thought to localize to the nuclear-facing side of the SPB (Kupke et al., 2011), so the Nbp1-Mps2 interaction we observe by FRET and BiFC could occur at the toroid region rather than the cytoplasm. The idea that the N-terminus of Mps3 is involved in SPB insertion is supported by analysis of NE fenestration in fission yeast, which implicates Sad1 phosphorylation as a trigger for NE remodeling (Fernandez-Alvarez et al., 2016). While putative Mps3 N-terminal phosphorylation sites (serine 70 and tyrosine 72) are non-essential (Fig 5A) (Li et al., 2017),deletion of the Mps3 N-terminus (*mps3Δ2-150*) exacerbates the growth defect of *mps2-381* or *mps3-F592S* (a mutation in a conserved SUN domain residue) (Fig 5B), providing evidence in budding yeast that the N- and C-terminus of Mps3 act cooperatively in SPB function.

**Figure 5.**
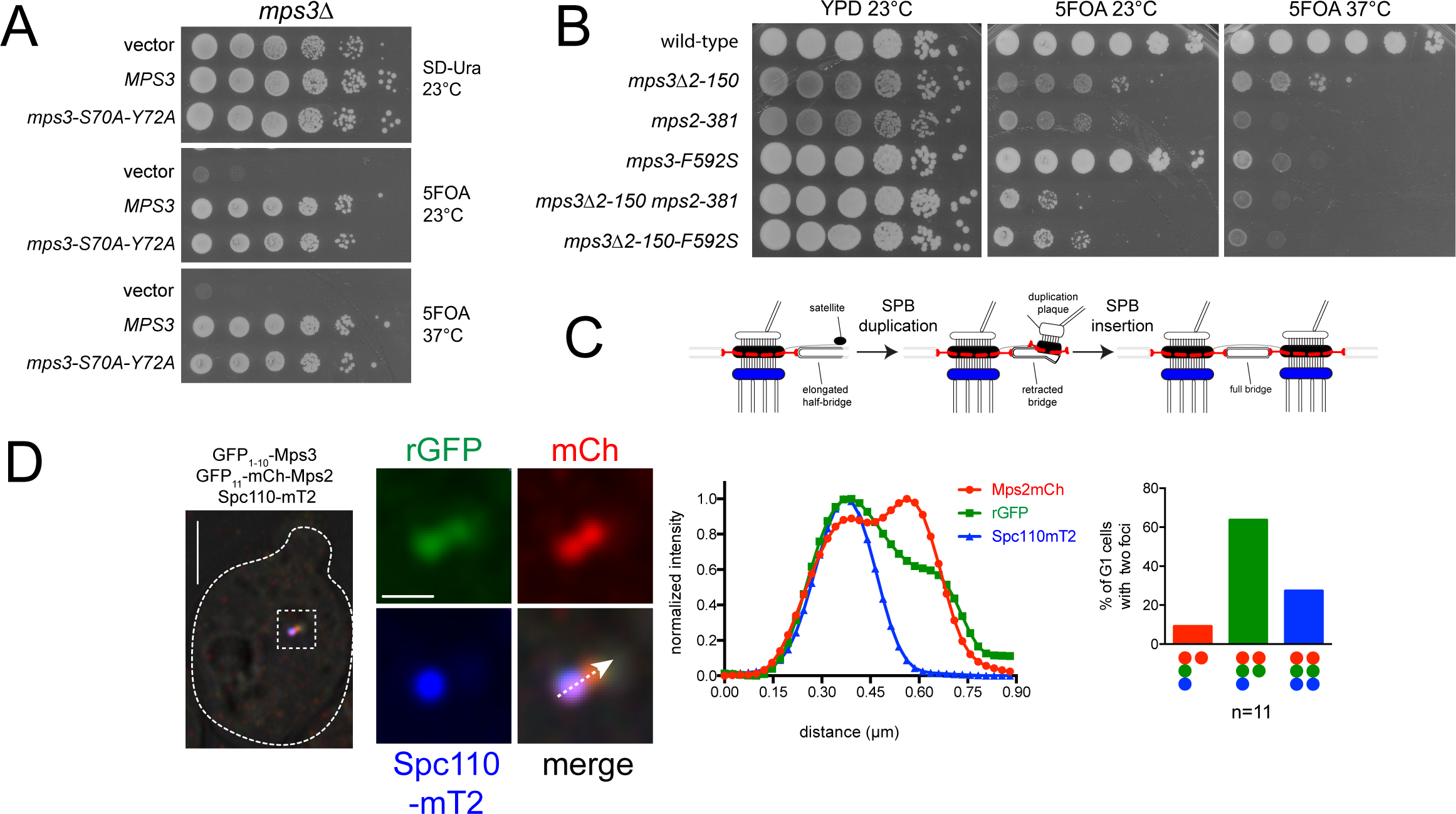
Role of the Mps3 N-terminus in SPB duplication. A. Putative phosphorylation sites at residues serine 70 and tyrosine 72 in the Mps3 N-terminus were mutated to alanine and tested for their ability to rescue growth of *mps3Δ pURA3-MPS3* (SLJ1053) in a serial dilution assay. Growth was compared to an empty vector or wild-type *MPS3* on 5-FOA, which selects for cells that have lost *pURA3-MPS3*. As a control, cells were also stamped to SD-URA. B. Growth of the indicated single and double mutants was compared to wild-type on 5-FOA at 23°C and 37°C. As a control, cells were also stamped to YPD at 23°C. Plates were incubated for 2 d at 37°C and for 3 d at 23°C. C. Schematic showing SPB duplication and NE insertion. Incorporation of Spc110 (blue) onto the new SPB requires its insertion into the NE, whereas Mps2 (red) localizes to the new SPB early in duplication before insertion (Burns et al., 2015; Pereira et al., 1999). D. Three color imaging of asynchronously growing cells (SLJ12884) expressing GFP_11_-mCherry-Mps2 (red) together with GFP_1-10_-Mps3 to assay binding by rGFP (green) relative to SPB insertion, determined by Spc110-mT2 incorporation (blue). A linescan across the magnified SPB region from a G1 cell undergoing SPB duplication illustrates that the rGFP is detected at the distal SPB before Spc110. 30 G1 cells were imaged, 11 of which had two Mps2 foci. The percentage of cells with two Mps2 foci that contained one or two foci of rGFP and Spc110 was quantitated. All observed combinations of SPBs with two foci are plotted. Bars, 2 µm (left panel) and 200 nm (center panels).

One prediction of the model that a non-canonical LINC complex formation drives membrane remodeling or stabilization is that binding of Mps2 and Mps3 N-termini at the new SPB occurs prior to, or during NE insertion (Fig 5C). To test this, we examined Spc110-mT2 in GFP_11_-mCherry-Mps2/GFP_1-10_-Mps3 containing cells to determine if rGFP appeared at the new SPB before or after Spc110 assembly. In no cases did we observe two foci of Spc110 before rGFP (Fig 5D). Thus, Mps2 and Mps3 N-termini interact before Spc110 is assembled, supporting the model that non-canonical LINC complex formation is involved in NE insertion.

In summary, the distribution of Mps3 and Bbp1 and pattern of Mps2 binding at SPB fenestra was unanticipated, highlighting the importance of using high resolution imaging to understand the structure of protein complexes *in vivo*. Our data calls into question the idea that a Bbp1-containing complex encircles the SPB and anchors the SPB core in the NE (Araki et al., 2010; Kupke et al., 2017; Schramm et al., 2000). An important avenue for future research is to understand the nature of additional NE tethering mechanisms at the SPB. One new candidate is Mps3, which we show is part of SPIN. Our results show that Mps2-Mps3 binding does not tether the bridge to the SPB (Jaspersen et al., 2006), but instead points to a novel role of SUN proteins in INM-ONM fusion and/or stabilization. Given the localization of SUN proteins at NPCs in mammalian cells and their role in de novo NPC assembly, which requires NE fenestration (Doucet and Hetzer, 2010; Funakoshi et al., 2011; Talamas and Hetzer, 2011), it will be interesting to examine if Sun1 also folds back on itself during NPC assembly. The luminal linkage of LINC complexes could drive INM and ONM approximation and the closed hairpin might stabilize the highly curved membrane that exists at NE fenestrae directly or through recruitment of other factors.

## Materials and Methods

### Yeast strains

Yeast strains are derivatives of W303 and are listed in Table S1. Standard conditions were used for yeast growth (Dunham et al., 2015). Deletion and tagging of genes was done using PCR-based methods in SLJ1070 (*Mata/Matα bar1/bar1 ade2-1/ADE2 trp1-1/TRP1 lys2Δ/LYS2 leu2-3,112/leu2-3,112 his3-11,15/his3-11,15 ura3-1/ura3-1*) and was verified by PCR (Gardner and Jaspersen, 2014; Longtine et al., 1998; Sheff and Thorn, 2004). Strains were made homozygous by tetrad dissection followed by crosses to generate diploids. In some cases, diploids arose spontaneously presumably because tagged combinations of SPB components or the deletion resulted in a mild defect in some aspect of spindle formation. The ploidy of all strains was verified by flow cytometric analysis of DNA content at the time of strain construction and when strains were grown for imaging experiments.

Construction of strains containing *mps2-381* or *mps3Δ2-150*/*mps3-F592S* has been previously reported (Bupp et al., 2007; Jaspersen et al., 2006). Double mutants were created using standard genetic methods (Dunham et al., 2015). pSJ546 (pRS314-*mps3*-S70A Y72A) was created by oligonucleotide-directed mutagenesis of pSJ154 (pRS314-*MPS3*) and was transformed into *mps3Δ::HIS3MX pURA3-MPS3* (SLJ1053).

pSJ2165 (pRS315-*NOP1pr-GFP*_*11*_*-mCh-MPS2*) was created by cloning a PCR fragment containing the *MPS2* ORF into the NheI and SalI sites of pSJ1321 (pRS315-*NOP1pr-GFP*_*11*_*-mCh-PUS1*) (Smoyer et al., 2016). A diploid strain containing the *MPS2* or *PUS1* reporter was constructed by transformation into indicated GFP_1-10_ tagged strains. N- and C-terminal tagging constructs for split-GFP (Smoyer et al., 2016) were used to create fusions to Mps3 and Heh2 by PCR in these strains. Haploid cells containing both halves of split-GFP were generated by sporulation followed by tetrad dissection.

### Yeast two-hybrid interactions

Strains SLJ1644 (wide-type), SLJ12623 (*pom152Δ*), SLJ12624 (*pom152Δ mps2Δ*) were co-transformed with binding- and activation-domain (BD and AD) fused plasmids, pOBD-Bbp1 (pSJ403), pOAD-Ndc1 (pSJ383) and pOAD-Spc29 (pSJ1828), respectively. Transformants were selected on SC-Leu-Trp plates, replica-plated to SC-Leu-Trp-His plates containing 25mM 3-amino-triazole (Sigma Aldrich) and incubated at 30°C for 4 d to detect interactions.

### SIM imaging

Cells were grown to an OD_600_ of 0.5-0.8 in freshly prepared imaging media (6.7 g Yeast-nitrogen base with ammonium sulfate without amino acids, 5 g casamino acids, 16.6 mg uracil and 950 ml ddH_2_O; after autoclaving, 4 ml 4 mg/ml adenine, 2 ml 4 mg/ml tryptophan and 50 ml 40% (w/v) sterile glucose added). Cells were fixed for 15 min in 4% paraformaldehyde (Ted Pella) in 100 mM sucrose, then washed two times in phosphate-buffered saline, pH 7.4. An aliquot of cells was resuspended in Dako mounting media (Agilent Technologies), placed on a cleaned glass slide, covered with a number 1.5 coverslip then allowed to cure overnight at room temperature.

SIM images were acquired with an Applied Precision OMX BLAZE (GE Healthcare) equipped with a 60X 1.42 NA Plan Apo oil objective. Images were collected in sequential mode with two or three PCO Edge sCMOS cameras (Kelheim, Germany) for each acquisition channel. Color alignment from different cameras in the radial plane was performed using the color alignment slide from GE Healthcare. In the axial direction, color alignment was performed using 100 nm TetraSpeck beads (ThermoFisher). Reconstruction was accomplished with the Softworx software according to manufacturer’s recommendations with a Wiener filter of 0.001. In most cases images are mT2/YFP with 514 nm excitation for YFP and then 445 nm excitation for mT2. In some cases, we modified the protocol for mT2/YFP/mCherry acquisition with the mCherry acquired first and excited with the 568 nm laser. The dichroic in every case was 445/514/561 with emission filters at 460-485 nm, 530-552 nm and 590-628 nm for mT2, YFP and mCherry, respectively. For image preparation, the SIM reconstructed images were scaled 4x4 and a max projection in z over the relevant slices was done.

### SPA-SIM

All single particle averaging was performed using custom macros and plugins for the open source program, ImageJ (NIH, Bethesda, MD). Plugins and source code are available for download at http://research.stowers.org/imagejplugins. Toroid alignment is fundamentally different than multi-spot alignment we previously described (Bestul et al., 2017; Burns et al., 2015) and therefore requires a different fitting strategy. For toroid alignment, we exclusively used Ndc1-YFP as our fiducial marker except in split-GFP experiments, where we used mCh signal associated with Mps2. Given the often incomplete appearance of toroids from 3D SIM microscopy (Fig S1A), we need a method to globally fit multiple parts of the ring simultaneously to a model that accounts 3D positioning of the ring as well as its rotation around the z axis of the microscope (ϕ) and its tilt with respect to that axis (θ).

We begin with the mathematical description of the ring itself. The following equation describes the travel in Cartesian space (x, y, z) around a ring of radius, r, which is tilted from the z axis by angle θ and then rotated about the z axis by angle ϕ:

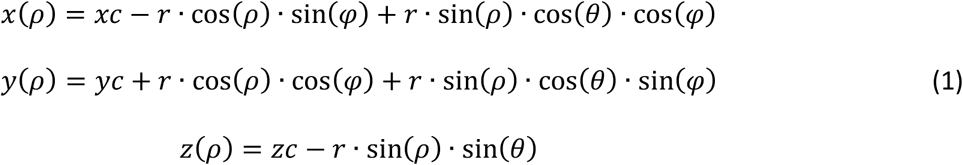

Here, xc, yc, and zc are the center of the ring in three dimensions and ρ is the angle that has been traveled about the ring.

Experimentally, a robust way to determine the positioning and orientation of a ring is to examine its xz cross section from its approximate center at multiple angles (in our case, we used 0, 45, 90, and 135 degrees, see Fig S1A). Because of the asymmetric resolution of the microscope, each point where the ring crosses is an asymmetric Gaussian function whose lateral dimension is the approximate lateral resolution of the microscope and whose vertical dimension is the approximate axial (z) resolution of the microscope. In order to improve the statistical accuracy of our cross sections, we average over a 2 pixel wide region for the xz cross sections.

Our task is now to fit a set of 8 cross sectional spots the tilted ring model described above. If we treat the initial guess of the center position of the ring as the origin, we must simply find the *p* values (and therefore 3D coordinates) at which the ring crosses the xz plane, the yz plane and the diagonal planes at 45 and 135 degrees. This is done by solving Eq. 1 for x = 0 (xz plane crossings), y = 0 (yz plane crossings), and x = y (45 degree crossings) and x = -y (135 degree crossings). The solutions were found with the aid of the Mathematica (Wolfram, Champaign, IL, USA) program as follows:

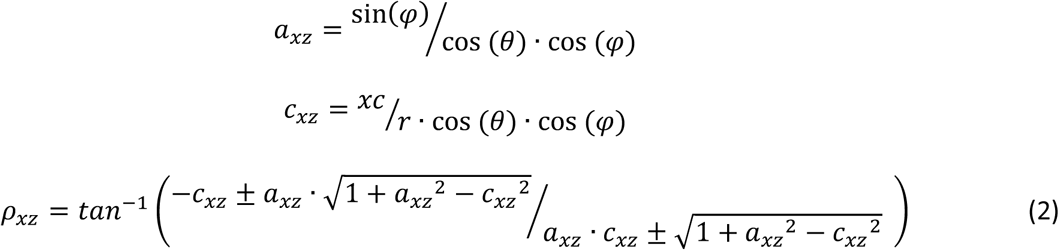

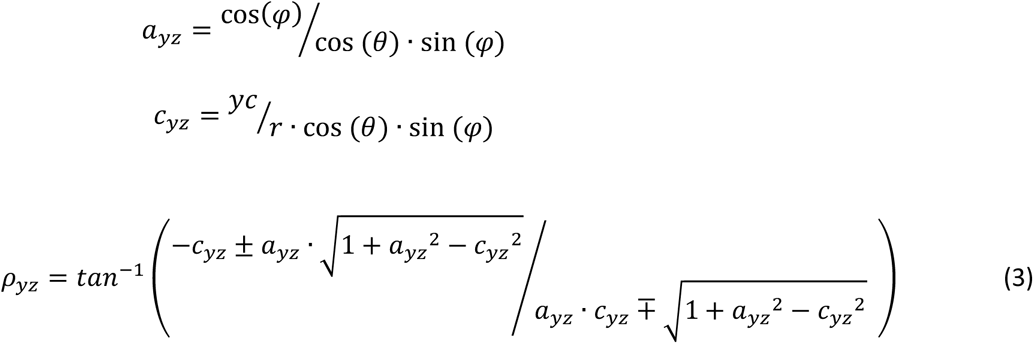

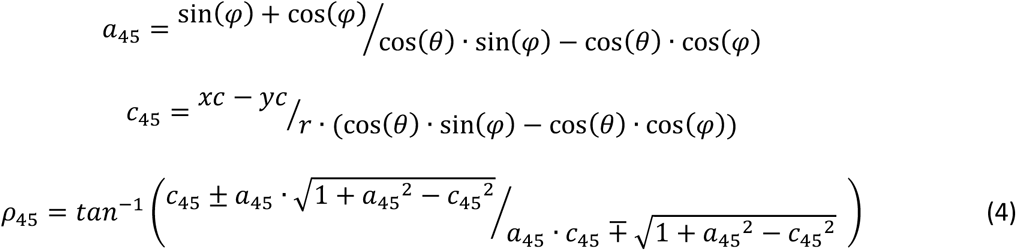

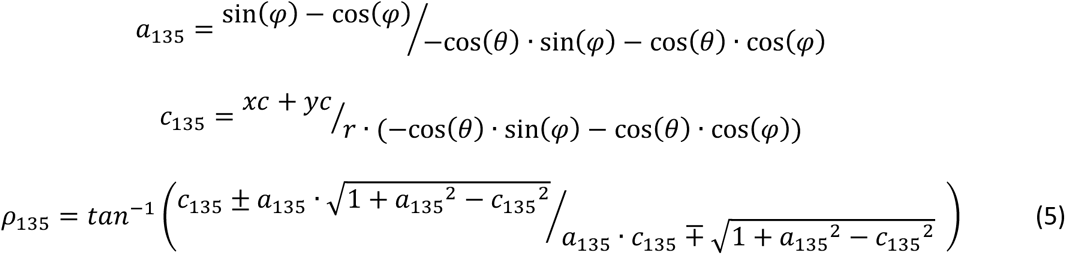

Care was taken to ensure that ρ values were between 0 and 2π and that the crossing points were not swapped. The final fit was a non-linear least squares global fit (Bevington and Robinson, 2003) to 8 asymmetric two dimensional gaussians. The standard deviations of these gaussians in x and z were linked together and the amplitudes on either side of each cross section were constrained to be no more than a factor of 2 different from one another. Four points were manually placed on the image at the approximate locations of the 0, 90, 180, and 270 degree crossing points of the ring. The radius was initialized from the average of the two distances derived from these points and the center in x and y was initialed at the center of these four points. The center in z was initialized at the maximum intensity position of the average of all of the z intensity profiles at these four points. The center of the toroid was constrained to be within 20% of the guess radius from the initialized center and the radius itself was constrained to be within a factor of 2 of its original initialized value. The z position was constrained to be within one z slice from its initialized position. The tilt angle (θ) was constrained to values less than 45 degrees. At low tilt angle values (for essentially flat rings), the rotation angle (ϕ) is poorly defined. As a result, we fixed the rotation angle (ϕ) at 10 degree increments and repeated the fit for every possible value of ϕ to ensure the best fit.

Quality of fit was assessed by visual inspection of the fitted cross-sectional images in comparison to the final fit in simulated cross-sectional images. The original 3D image was then transformed so that the fitted toroid was flat at the center of the final transformed image. In some cases, the final images were randomly rotated about their centers (in the xy plane) to avoid accidental non-homogeneous regions in the aligned image. The final SPA-SIM image was formed by averaging the realigned images. In rare cases, one image was much brighter than the others in a series; when this occurred, the bright image was normalized by its ratio to the other image intensities to avoid that image dominating the averaged image.

In some cases, there is reason to believe that the distribution of the secondary (not fiducial) channel is oriented at a specific direction from the center of the toroid (e.g. Mps3, which is localized on the half-bridge (Jaspersen et al., 2002)). For these, the angle of each individual toroid’s rotation was determined by manual drawing of a line between the ring center and the secondary distribution center. Images were then rotated so that the asymmetry is either pointing upwards or sideways before averaging. These cases are specifically pointed out in the text.

Radial profiles were generated using custom software interpolating pixel values at 1 pixel arc lengths and averaging around ever-expanding circles from the center of each aligned averaged image. Diameters were determined by fitting intensity profiles drawn through the vertical and horizontal centers of the averaged ring images to two Gaussian functions. Errors were determined by Monte Carlo analysis as described in (Burns et al., 2015). In cases where toroids appear approximately symmetric, the reported diameters were the average of vertical and horizontal values with propagated errors. In asymmetric cases (e.g. Mps3), we independently report the vertical and horizontal diameters.

## FRET

Cells were grown and fixed identically to SIM samples. An aliquot of fixed cells was resuspended in ProLong Diamond Antifade mounting media (ThermoFisher Scientific), placed on a cleaned glass slide, covered with a number 1.5 coverslip then allowed to cure overnight at room temperature. Images were acquired on a Nikon Eclipse TI equipped with a Yokogawa CSU W1 spinning disk head and Andor EMCCD using a Nikon Apo TIRF 100X 1.49NA Oil objective. mTurquoise2 was imaged using a 445nm laser and 480/30 emission filter with a maximum power of 1.2mW measured at the sample. YFP was imaged using a 514nm laser and ET535/30m emission filter with a maximum power of 2.5mW measured at the sample. For each sample 16 points were manually or automatically selected depending on cell density. Afterwards, an automation script moved to positions, found focus using Nikon PFS, imaged mT2/YFP, bleached at 514nm for one minute and re-imaged. Image processing was performed in ImageJ using custom macros and plugins (https://github.com/jouyun/). In brief, a small blurring was performed, followed by a background subtraction and registration; puncta were identified using a local maximum finder and adaptive region grow; these were quantified in the mT2 channel before and after the bleach. Average FRET values with donor and acceptor and donor only were determined and statistical significance from the donor only control was determined using the Student’s t-test.

### Bimolecular fluorescence complementation assay

BiFC complementation was assayed in conjunction with SIM, as described above. For split-GFP, mCherry/GFP with 568 nm excitation for mCherry and then 488 nm excitation for GFP. The dichroic in every case was 568/448 with emission filters at 590-628 nm and 504-552 nm for mCherry and GFP, respectively. Note, the pixelated appearance of mCherry-GFP_11_-Pus1 in SIM images is due to the SIM reconstruction.

Samples for three color images were prepared as described for SIM, however, images with split-GFP, mCherry and mTurquoise2 were taken on a Zeiss-LSM780 equipped with an AiryScan super resolution add-on using a 63X 1.4NA Oil objective. mCherry was imaged first, using the 561 nm laser line and a 488/561 nm excitation dichroic; a LP575 nm emission filter was in front of the AiryScan detector. The mCherry signal was then photobleached with multiple rounds of excitation with 100% laser power, and a second mCherry image was acquired to ensure complete bleaching. This allowed us to then image split-GFP with 514 nm excitation and a 458/514 nm dichroic; a LP525 nm emission filter was in front of the detector. Lastly, mTurqouise2 was imaged with 458 nm excitation, and emission was collected through a 420-480 nm BP filter. We verified that no GFP signal was collected in this range. This scheme was needed to avoid cross-talk, as mCherry (but not mTurquoise2) is excited at 514 nm; the usual 488 nm excitation used for GFP will excite both split-GFP and mTurquoise2. Standard AiryScan settings were used for image acquisition. 21 z-slices were acquired with 0.22 µm spacing. A zoom of 2.5 was used, with 1104 x 1104 pixels per image, giving a pixel size, prior to processing, of 50 nm. Pixel dwell time was 2.92 µs. AiryScan super resolution processing was carried out using Carl Zeiss Zen software, with a Weiner filter setting of 7.5. For post-processing, images were color aligned in ImageJ to account for small motion during imaging and smoothed with a Gaussian blur of radius 1 pixel. Images were scaled 2 x 2 with bilinear interpolation. Far red beads (ThermoScientific) embedded in the mounting media next to the yeast were imaged at 633 nm and used for AiryScan super resolution alignment.

### Transmission electron microscopy

Cells were grown overnight at 30°C in YPD, arrested in G1 using 1 μg/ml α-factor, then grown for an additional hour in YPD at 30°C to enrich for cells undergoing duplication. Cells were quickly harvested and frozen on the Leica EM-Pact (Wetzlar, Germany) at ∼2050 bar, transferred under liquid nitrogen into 2% osmium tetroxide/0.1% uranyl acetate/acetone, and transferred to the Leica AFS. The freeze substitution protocol was as follows: -90° for 16 h, raised 4°/h for 7 h, -60° for 19 h, raised 4°/h for 10 h, and -20° for 20 h. Samples were then removed from the AFS, placed in the refrigerator for 4 h, and then allowed to incubate at room temperature for 1 h. Samples went through three changes of acetone over 1 h and were removed from the planchettes. They were embedded in acetone/Epon mixtures to final 100 % Epon over several days in a stepwise procedure as described (McDonald, 1999). Sixty-nanometer serial thin sections were cut on a Leica UC6, stained with uranyl acetate and Sato’s lead, and imaged on a FEI Tecnai Spirit (Hillsboro, OR).

## Acknowledgements

We are grateful to Jenny Friederichs for help with EM and to Joe Varberg, Briana Holt and Shelly Jones for comments on the manuscript. Research reported in this publication was supported by the Stowers Institute for Medical Research and the National Institute of General Medical Sciences of the National Institutes of Health under award number R01GM121443 (to SLJ). Original data underlying this manuscript can be downloaded from the Stowers Original Data Repository at http://www.stowers.org/research/publications/LIBPB-1349. The authors declare no competing financial interests.

## Author Contributions

SLJ, JC and JRU conceived the experiments, JC and JMG constructed strains and performed arrests, JC, JMG, BRS and ZU performed SIM, SES and SM performed and analyzed FRET, JRU developed tools for image analysis, SLJ prepared figures and wrote the paper with input from all the authors.

## Abbreviations

MTOC: microtubule-organizing center
SPB: spindle pole body
SIM: structured illumination microscopy
EM: electron microscopy
NE: nuclear envelope
NPC: nuclear pore complex
SPA: single particle analysis
SEM: standard error of the mean
EM: electron microscopy
SPIN: spindle pole body insertion network
SUN: Sad1-UNC-84 homology
KASH: Klarisht-ANC-1-Syne-1 homology
LINC: linker of nucleoskeleton and cytoskeleton
BiFC: bimolecular fluorescence complementation
5-FOA: 5-fluoro-orotic acid
3-AT: 3-amino-triazole
mT2: mTurquoise2

## Supplementary Material

**Figure S1.**
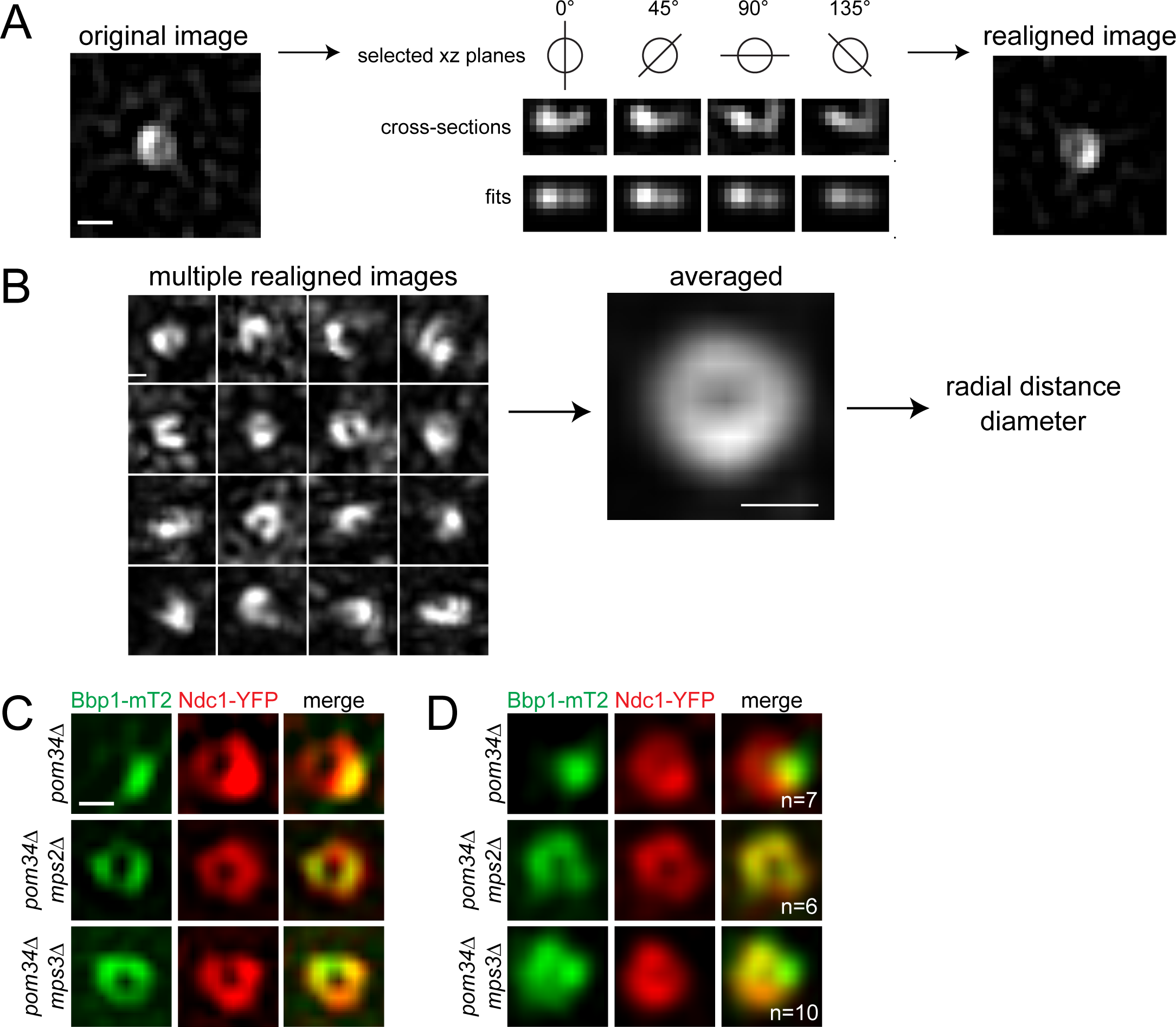
3D single particle averaging of toroidally distributed proteins. A. SIM of Ndc1-YFP, showing ring-like distribution. The cross-sectional view in selected xz planes is shown, along with fit values used to produce the realigned image. B. A montage of realigned Ndc1-YFP images are shown, which are then averaged to create a projection. From this, radial intensity profiles from the center of the SPB outward can be determined and used to estimate diameter. C-D. Localization of Ndc1-YFP (red) and Bbp1-mT2 (green) in individual SIM images (C) or averaged images (D) from *pom34Δ* (SLJ12300), *pom34Δ mps2Δ* (SLJ10936) and *pom34Δ mps3Δ* (SLJ11913). Number of SPBs included in average is shown. Bars in Fig S1, 100 nm.

**Figure S2.**
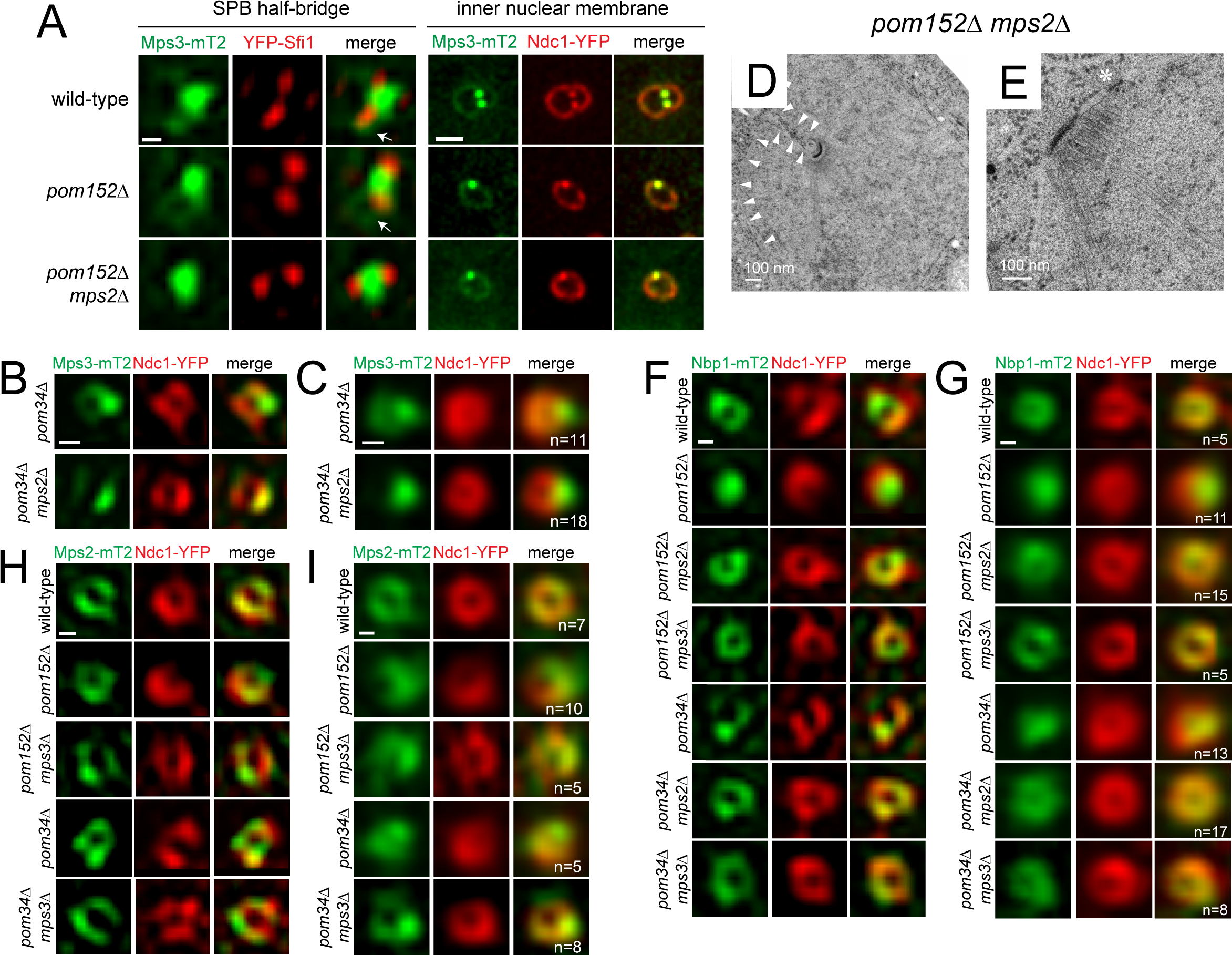
Loss of Mps2 specifically disrupts Mps3 at the toroid. A. SIM showing localization of YFP-Sfi1 (red) and Mps3-mT2 (green) in G1 cells selected from an asynchronously growing population of wild-type (SLJ12060), *pom152Δ* (SLJ12100) *or pom152Δ mps2Δ* (SLJ12100) cells. Arrows point to Mps3-mT2 that can be seen around one YFP-Sfi1 focus, which is likely to be the mother SPB, in wild-type and *pom152Δ* but not *pom152Δ mps2Δ* cells. Localization of Ndc1-YFP (red) and Mps3-mT2 (green) in G1 cells selected from an asynchronously growing population of wild-type (SLJ10636), *pom152Δ* (SLJ11071) and *pom152Δ mps2Δ* (SLJ10535). Bar, 100 nm (left panel), 2 µm (right panel). B-C. Individual SIM (B) and averaged (C) images showing localization of Ndc1-YFP (red) and the distribution of Mps3-mT2 (green) in *pom34Δ* (SLJ11165), *pom34Δ mps2Δ* (SLJ10899). Bars, 100 nm. D-E. EM from *pom152Δ mps2Δ* (SLJ4270) showing an intact SPB on a NE invagination. Small arrowheads show the location of the NE. In E, an SPB is embedded at NE with an NPC nearby. The asterisk marks the NPC. Bars, shown. F-G. Localization of Ndc1-YFP (red) and Nbp1-mT2 (green) individual SIM images (F) and averaged images (G) from wild-type (SLJ10898), *pom152Δ* (SLJ12301), *pom152Δ mps2Δ* (SLJ10863), *pom152Δ mps3Δ* (SLJ10864), *pom34Δ* (SLJ11970), *pom34Δ mps2Δ* (SLJ10900) and *pom34Δ mps3Δ* (SLJ11969) strains. Number of SPBs included in average is shown. H-I. Localization of Ndc1-YFP (red) and Mps2-mT2 (green) in individual SIM images (H) and averaged images (I) from wild-type (SLJ11171), *pom152Δ* (SLJ12370), *pom152Δ mps3Δ* (SLJ11173), *pom34Δ* (SLJ12303) and *pom34Δ mps3Δ* (SLJ11968) strains. Number of SPBs included in average is shown. Bars in Fig S2F-I, 100 nm.

**Table S1. Yeast Strains.**

**See attached table**

\# All strains are *ADE2* derivatives of W303, unless indicated.

* Denotes strains used for the yeast two-hybrid assay that are: *trp1-901 leu2-3,112 ura3-52 his3-200 gal4Δ gal80Δ LYS2::GAL1-HIS3 met2::GAL7-LACZ GAL1-ADE2*

** Denotes strains derived from the BY strain background that are: *can1Δ::STE2pr-HIS3MX lyp1Δ his3Δ1 leu2Δ0 ura3Δ0 met15 Δ0*

